# The H_abc_ Domain of Syntaxin 3 is a Ubiquitin Binding Domain with Specificity for K63-Linked Poly-Ubiquitin Chains

**DOI:** 10.1101/219642

**Authors:** Adrian J. Giovannone, Elena Reales, Pallavi Bhattaram, Sirpi Nackeeran, Adam B. Monahan, Rashid Syed, Thomas Weimbs

## Abstract

Syntaxins are a family of membrane-anchored SNARE proteins that are essential components required for membrane fusion in all eukaryotic intracellular membrane trafficking pathways. Syntaxins contain an N-terminal regulatory domain, termed the H_abc_ domain, that is not highly conserved at the primary sequence level but folds into a three-helix bundle that is structurally conserved among family members. The syntaxin H_abc_ domain has previously been found to be structurally very similar to the GAT domain present in GGA family members and related proteins that are otherwise completely unrelated to syntaxins. Because the GAT domain has been found to be a ubiquitin binding domain we hypothesized that the H_abc_ domain of syntaxins may also bind to ubiquitin. Here, we report that the H_abc_ domain of syntaxin 3 (Stx3) indeed binds to monomeric ubiquitin with low affinity. This domain binds efficiently to K63-linked poly-ubiquitin chains within a narrow range of chain lengths but not to K48-linked poly-ubiquitin chains. Other syntaxin family members also bind to K63-linked polyubiquitin chains but with different chain length specificities. Molecular modeling suggests that residues of the GGA3-GAT domain known to be important for ionic and hydrophobic interactions with ubiquitin have equivalent, conserved residues within the H_abc_ domain of Stx3. We conclude that the syntaxin H_abc_ domain and the GAT domain are both structurally and functionally related, and likely share a common ancestry despite sequence divergence. Binding of K63-linked poly-ubiquitin chains to the H_abc_ domain may regulate the function of syntaxins in membrane fusion or may suggest additional functions of this protein family.

## Introduction

SNARE (soluble *N*-ethylmaleimide-sensitive factor attachment protein receptor) proteins are the indispensable mediators of membrane fusion reactions within the endomembrane system of eukaryotic cells [1-5]. The SNARE superfamily consists of several sub-families whose members contain one or two SNARE domains of ∽60 residue length [3, 4, 6]. Members of the syntaxin family of SNAREs are central to the formation of SNARE complexes. They contain a C-terminal transmembrane anchor, preceded by the SNARE domain. The latter engages in interactions with cognate SNAREs to form a SNARE complex in a 4-helix bundle arrangement that ultimately leads to membrane fusion [7-9].

Syntaxins also contain an N-terminal regulatory domain that consists of three a helices (a,b,c) and has been termed the H_abc_ domain. At least 16 syntaxins are encoded in the human genome and many more in divergent species. Amongst these syntaxins, the H_abc_ domains are poorly – or not at all - conserved at the primary sequence level. However, in the cases of syntaxin family members whose H_abc_ domains have been structurally studied it was found that they all share a highly conserved fold. This includes Stx1A [10, 11], Stx6 [12], Sso1 [13], Stx10 [14], Vam3p [15] and several others (see Protein Data Bank) whose H_abc_ domains fold into essentially superimposable three-helix bundles despite limited or absent sequence similarity. The H_abc_ domains of various syntaxins have been found to be binding sites to proteins that regulate SNARE function including those of the munc18, munc13 and synaptotagmin families [16]. In addition, in some – but not all – syntaxins the H_abc_ domains have the ability to engage in an intramolecular interaction with the SNARE domain resulting in a tetrameric helical bundle. This “closed” conformation generally inhibits the formation of complexes with cognate SNARE proteins and thereby inhibits membrane fusion [16, 17]. These findings clearly indicate that the H_abc_ domains of syntaxins play a critical role in the regulation of membrane fusion and that this function depends on the conserved three-dimensional structure of these domains.

It was surprising when it was found that a conserved domain in a very different family of proteins shares the same fold with the syntaxin H_abc_ domain. The GAT (GGAs and TOM) domain of GGA1 was found to be nearly superimposable with the H_abc_ domains of Stx1A and Stx6 despite their lack of sequence similarity [18]. GGA proteins are Golgi- and endosome-associated clathrin adaptor proteins involved in cargo recruitment in membrane trafficking pathways. At the time of the discovery of the structural similarity with the syntaxin H_abc_ domain, relatively little was known about the function of the GAT domain. Subsequently, however, the GAT domains of GGA proteins were found to be ubiquitin binding domains [19-24] which helped to explain their function in recruiting ubiquitinated membrane proteins for targeting to multivesicular bodies (MVBs) [25].

Ubiquitin, an 8 kDa protein, is covalently attached to lysine residues of target proteins via E3 ligases [26]. In the case of membrane proteins, reversible ubiquitination serves as a signal for targeting to endosomes, and then to intraluminal vesicles of MVBs. MVBs can subsequently either fuse with lysosomes leading to degradation [27] or they can fuse with the plasma membrane leading to extracellular secretion of membrane proteins in the form of exosomes [28]. Ubiquitin itself may be ubiquitinated at any of its seven lysine residues leading to target proteins being tagged with a chain of polyubiquitin molecules [29]. K48-linked polyubiquitin chains are a signal for proteasomal degradation whereas K63-linked polyubiquitin chains are another signal for trafficking of membrane proteins to the endosomal pathway, and especially into the MVB pathway [30-32]. The GAT domain of GGA proteins has been shown to be required for the sorting of membrane proteins tagged with K63-polyubiquitin chains into the MVB pathway [31, 32].

GGA proteins themselves are also mono-ubiquitinated in a manner dependent on the binding of ubiquitin to their GAT domain [19]. A large number of ubiquitin-binding proteins have been found to also be ubiquitinated themselves. The reasons for this are not always completely clear but it is thought that concurrent ubiquitin-binding and ubiquitination of sorting proteins aids in the establishment of protein networks to create sorting domains [25, 33].

We have recently reported that syntaxin 3, a SNARE involved in membrane fusion at the apical plasma membrane of polarized epithelial cells, undergoes mono-ubiquitination at lysine residues adjacent to its transmembrane domain [34]. Ubiquitination of Stx3 leads to endocytosis from the basolateral plasma membrane, direction into the endosomal/MVB pathway and eventually excretion with exosomes [34]. Functional studies using a non-ubiquitinatable Stx3 mutant suggested that Stx3 may function to sort specific cargo proteins into the MVB/exosomal pathway [34]. Such a function is unexpected for a protein thought to be involved in membrane fusion.

The structural similarity between the H_abc_ domain of syntaxins and the GAT domain suggests a common ancestry and related function. Given this structural similarity, and given the finding that Stx3 – like GGA proteins - is mono-ubiquitinated and appears to play a role in cargo sorting in the MVB/exosomal pathway, we hypothesized that the H_abc_ domain of Stx3 may be a ubiquitin-binding domain. We report here that Stx3 indeed binds to ubiquitin with low affinity, and with much higher affinity to K63-linked polyubiquitin chains, but not K48-linked polyubiquitin chains. Structural modeling and mutagenesis experiments suggest that the mode of ubiquitin binding is similar to that of the GAT domain and may involve conserved hydrophobic interactions and a salt bridge. These results suggest that syntaxin function may be regulated by polyubiquitin binding, and that syntaxins may function in protein sorting in addition to their established role in membrane fusion.

## Results

### Stx3 binds non-covalently to ubiquitin

To investigate the possibility that Stx3 may bind to ubiquitin we incubated a purified GST-fusion protein of the entire cytoplasmic domain (1-265) of Stx3 with ubiquitin-coated beads to observe any interaction in a pull-down assay. A GST-fusion protein of the GGA3-GAT domain served as a positive control. As shown in Fig. 1A, GGA3-GAT interacts strongly with ubiquitin-coated beads as expected. GST-Stx3 also interacts with ubiquitin-coated beads although with reduced efficiency as compared to GGA3-GAT. A GST-fusion protein containing only the H_abc_ domain (1-146) of Stx3 pulls down with ubiquitin-coated beads much more efficiently than the entire Stx3 cytoplasmic domain (Fig. 1B). This suggests that the H_abc_ domain directly binds to ubiquitin, and that this interaction is inhibited by intramolecular binding between the H_abc_ and SNARE domains in the “closed conformation” of Stx3. To test this possibility, we introduced the LE165/166AA mutation that has previously been shown to prevent the closed conformation in the highly conserved Stx1A leading to a constitutively open conformation [17]. The observed increase in binding of the open-mutant vs. wild-type Stx3 (Fig. 1B) suggests that the H_abc_ domain preferentially binds to ubiquitin when it is not engaged in binding to the SNARE domain.

**Figure 1.**
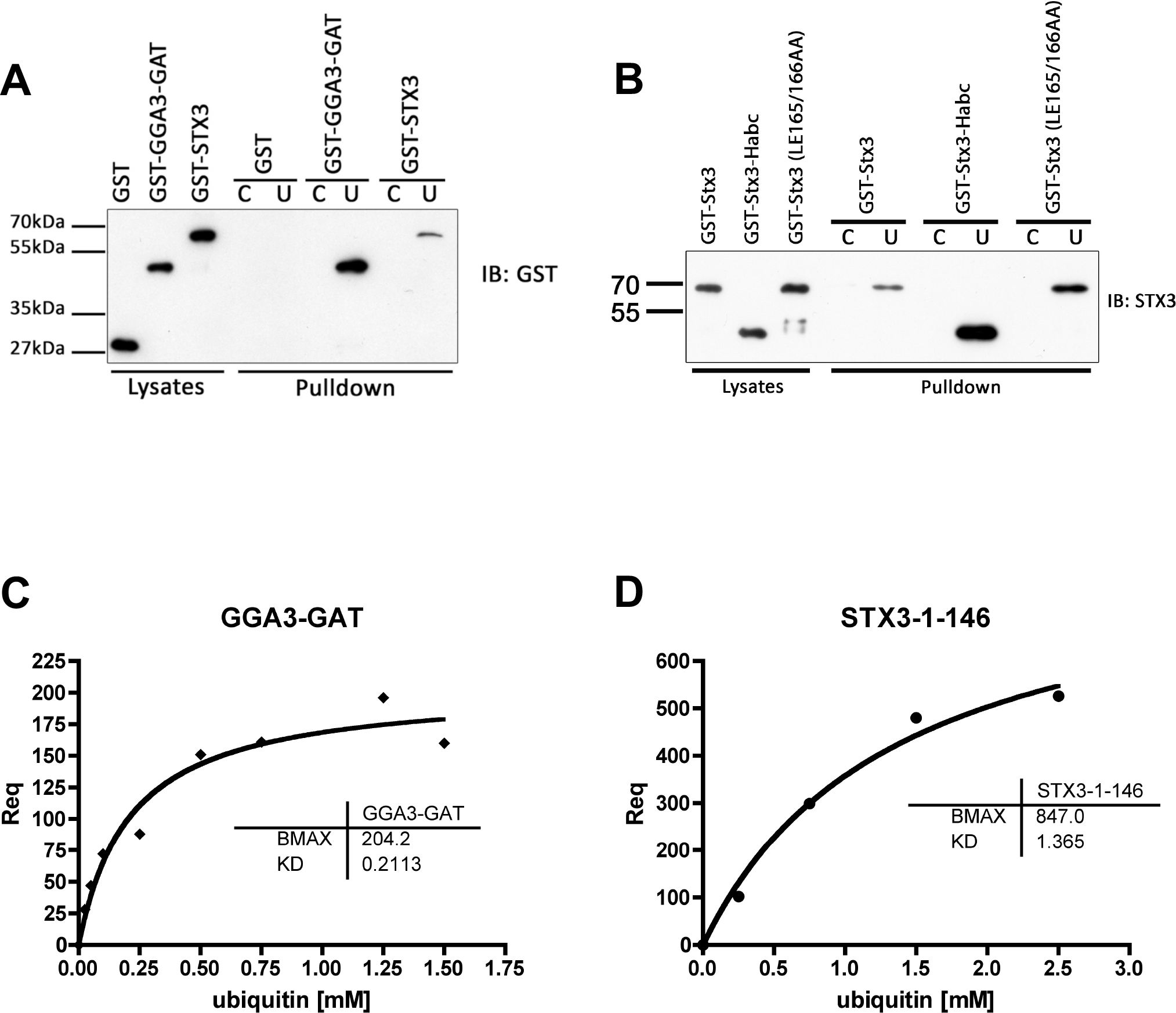
Stx3 binding to mono-ubiquitin. **(A)** Purified GST fusion protein of the GGA3-GAT domain (positive control for ubiquitin binding) or the cytoplasmic region (1-265) of Stx3 was precipitated with ubiquitin-coated (U) agarose beads or control, uncoated CL4B beads (C) and subjected to immunoblotting using anti-GST antibody. **(B)** Purified GST fusion proteins: cytoplasmic region of Stx3 (1-265), H_abc_ domain Stx3 (1-146), and constitutively open mutant of cytoplasmic region Stx3 (L165A/E166A) were each precipitated as in panel A and probed with anti-Stx3 antibody. **(C)** SPR experimental data of interaction between captured GST-GGA3-GAT and free ubiquitin. **(D)** SPR experimental data of interaction between captured GST-Stx3-1-146 and free ubiquitin.

To assess the affinity of the H_abc_ domain of Stx3 for ubiquitin, we utilized a surface plasmon resonance assay using immobilized GST-Stx3-H_abc_ in comparison with GST-GGA3-GAT as a positive control. The measured K_d_ for the binding of GST-GGA3-GAT to ubiquitin is 0.211 mM which is consistent with previously published values of 0.231 mM [24] and 0.181 mM [23]. This interaction has been described as a “high-affinity” interaction for a ubiquitin binding protein [29]. In comparison, the measured K_d_ for the H_abc_ domain of Stx3 is approximately four times weaker (1.36 mM) (Fig. 1D) indicating that the binding of Stx3 to mono-ubiquitin in solution is a low-affinity interaction.

Altogether, these results suggest that the similarity between the GAT domain of GGA proteins and the H_abc_ region of Stx3 is not only structural, but also functional with respect to ubiquitin binding.

### Stx3 binds to K63-linked but not K48-linked polyubiquitin chains

Ubiquitin-binding domains commonly have weak affinities for mono-ubiquitin but the presence of multiple ubiquitin binding sites in the same molecule often results in much higher affinities for polyubiquitin chains [29]. Since the affinity of the H_abc_ domain of Stx3 for mono-ubiquitin was low, we next investigated whether Stx3 may exhibit higher affinity for polyubiquitin chains. The two predominant chain-linkages are via the K48 or K63 residues of ubiquitin, respectively. GST-Stx3 was incubated with either K48- or K63-linked polyubiquitin chains covering a range of lengths from monomeric to 7-mers. As shown in Fig. 2A, Stx3 interacts with K63-linked polyubiquitin chains of lengths between 3-5. In this assay, no binding is detected to monomeric ubiquitin consistent with the weak affinity of Stx3 to monomeric ubiquitin measured above (Fig. 1D). Stx3 also does not exhibit interaction with K63-linked ubiquitin chains longer than 5 units suggesting selectivity to a narrow range of chain lengths. Importantly, no interaction between Stx3 and K48-linked chains of any length is detected suggesting that Stx3 shows strong preference to the K63 linkage.

**Figure 2.**
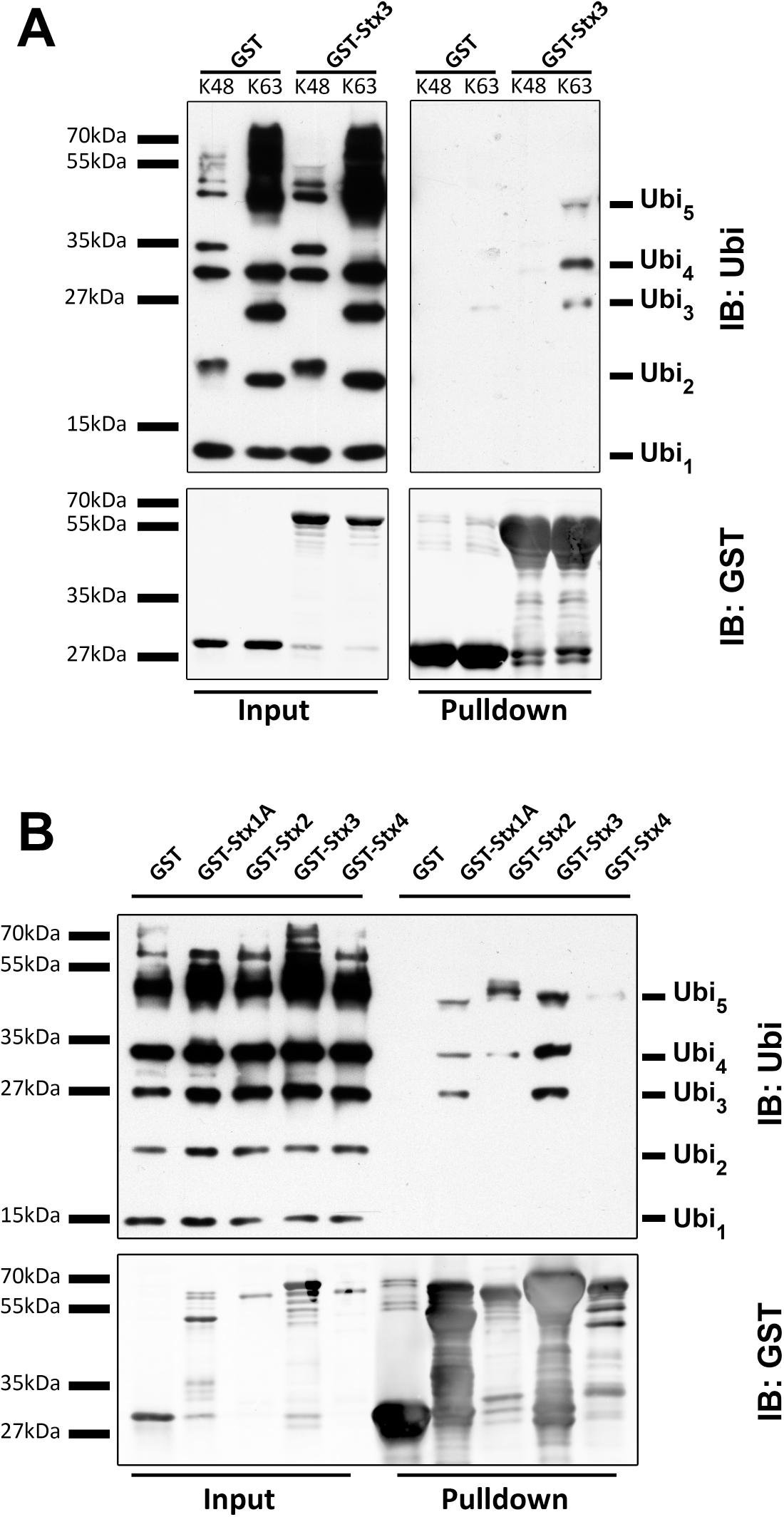
Stx3 binding to K63-linked polyubiquitin chains. **(A)** Purified GST fusion protein of Stx3 (cytoplasmic region, 1-265) was incubated with a mix of K48 or K63-linked polyubiquitin chains (1-7 ubiquitin molecules in length) followed by pull-down with glutathione sepharose and immunoblot (IB) using anti-ubiquitin or anti-GST antibodies. **(B)** Purified GST fusion protein of the cytoplasmic regions of Stx1A, Stx2, Stx3, or Stx4 were incubated with a mix of K63-linked polyubiquitin chains (1-7 ubiquitin molecules in length) followed by glutathione sepharose pull-down and IB using anti-ubiquitin and anti-GST antibodies.

Next, we tested whether the ability to interact with K63-linked ubiquitin chains is unique to Stx3 or may also be a feature of other syntaxins. We compared GST-fusion proteins of Stx1A, Stx2, and Stx4 side-by-side with Stx3. Stx1A and Stx2 interact with K63-linked ubiquitin chains similarly to Stx3 albeit with somewhat differing preferences for different chain lengths (Fig. 2B). Stx4 only exhibits very weak interaction with Ubi_5_. None of these syntaxins show binding to monomeric ubiquitin in this assay. Altogether, these data suggest that the H_abc_ domains of several syntaxins are capable of binding to ubiquitin chains and that Stx3 shows strong preference for K63-linked ubiquitin chains of a narrow range of lengths.

### Structural modeling

The GAT domain of GGA3 has been to shown have two distinct binding sites for ubiquitin. Site 1 has been studied in most detail and encompasses residues from the C-terminal half of helix A and the N-terminal half of helix B (Fig. 3A). X-ray structure analysis of ubiquitin in complex with the GGA3-GAT domain revealed prominent hydrophobic interactions with ubiquitin involving L227, M231 and L247 of GGA3, and a salt bridge involving E246 and E250 of GGA3 [23, 24]. These residues interact closely with I44 and R42, respectively, of ubiquitin. The other ubiquitin binding site of GGA3 (site 2) is located on the opposite face of the 3-helix bundle of the GAT domain and encompasses residues in helices B and C [24], but no 3D structure is available for this interaction.

**Figure 3.**
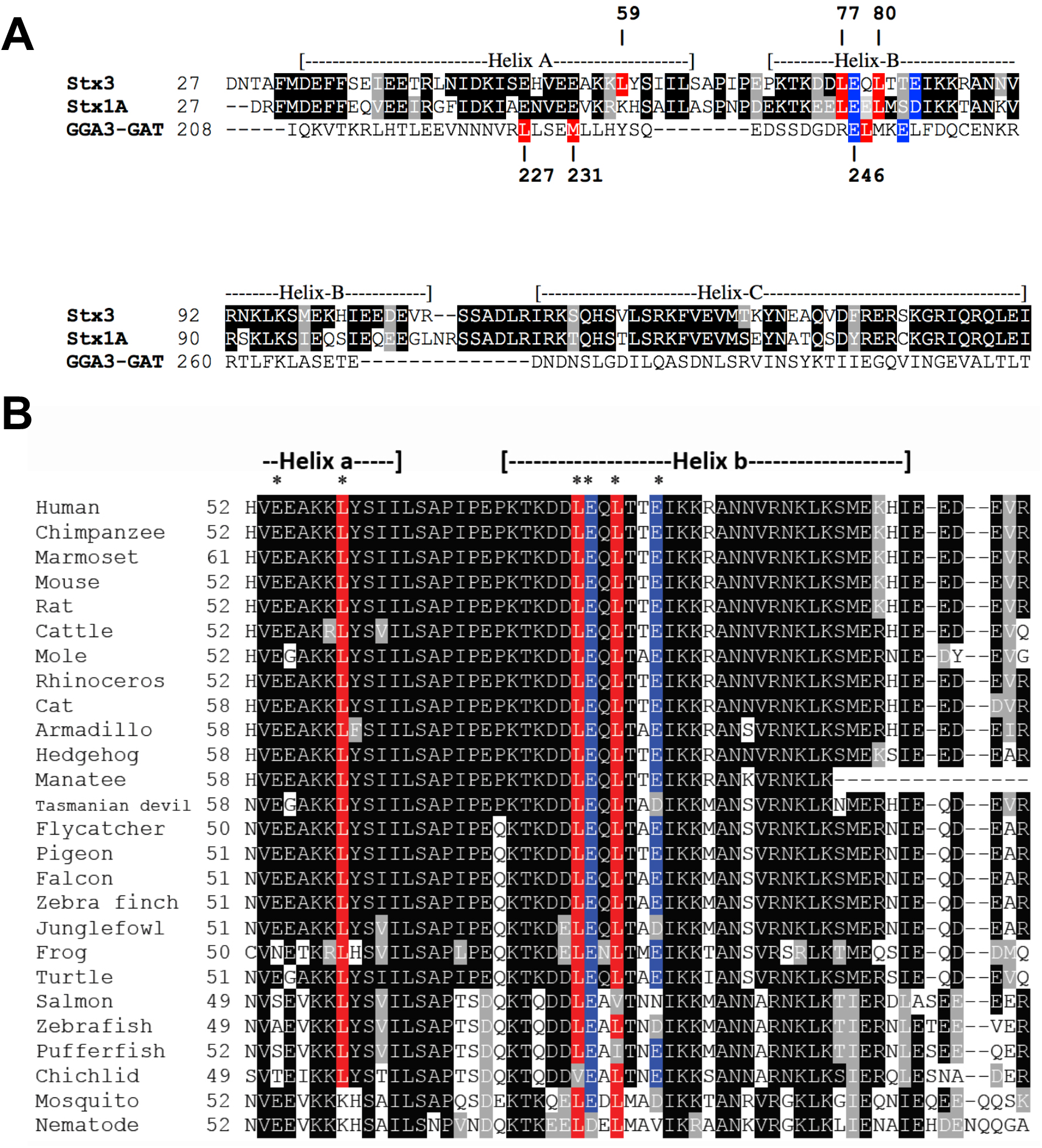
Conservation of residues important in GGA3-ubiquitin interaction. **(A)** Sequence alignment of the H_abc_ domains of human Stx3 and human Stx1A with the GAT domain of human GGA3. Hydrophobic and ionic residues known to be involved in the interaction between GGA3 and ubiquitin are highlighted in color. Putative equivalent residues, determined based on structural alignment (Fig. 4) in syntaxins are similarly highlighted. **(B)** Sequence alignment of Stx3 orthologues from 26 different species. Hydrophobic and ionic residues that are putatively involved in ubiquitin interaction are highlighted in red and blue, respectively.

To better understand how the H_abc_ domain of Stx3 may interact with ubiquitin we constructed a model based on the X-ray structure of ubiquitin in association with site 1 of the GGA3 GAT domain [23, 24]. We modeled the Stx3 sequence into the known structure of the Habc domain of the closely related Stx1A [10, 11], and then fitted this Stx3 H_abc_ domain onto the GGA3 GAT domain. The resulting model is shown in Fig. 4. This allowed us to identify Stx3 residues that correspond most closely to the known hydrophobic and ionic interactions between GGA3-GAT and ubiquitin. As shown in Fig. 4, the Stx3 H_abc_ domain has two glutamic acid residues (E78 and E83) in very similar positions as E246 and E250 of GGA3-GAT, and these residues maybe be predicted to engage in a salt bridge with R42 of ubiquitin. Similarly, L59, L77 and L80 of Stx3 would form a hydrophobic pocket that may interact with I44 of ubiquitin, analogous to the hydrophobic pocket formed by L227, M231 and L247 of GGA3 (Fig. 4).

**Figure 4.**
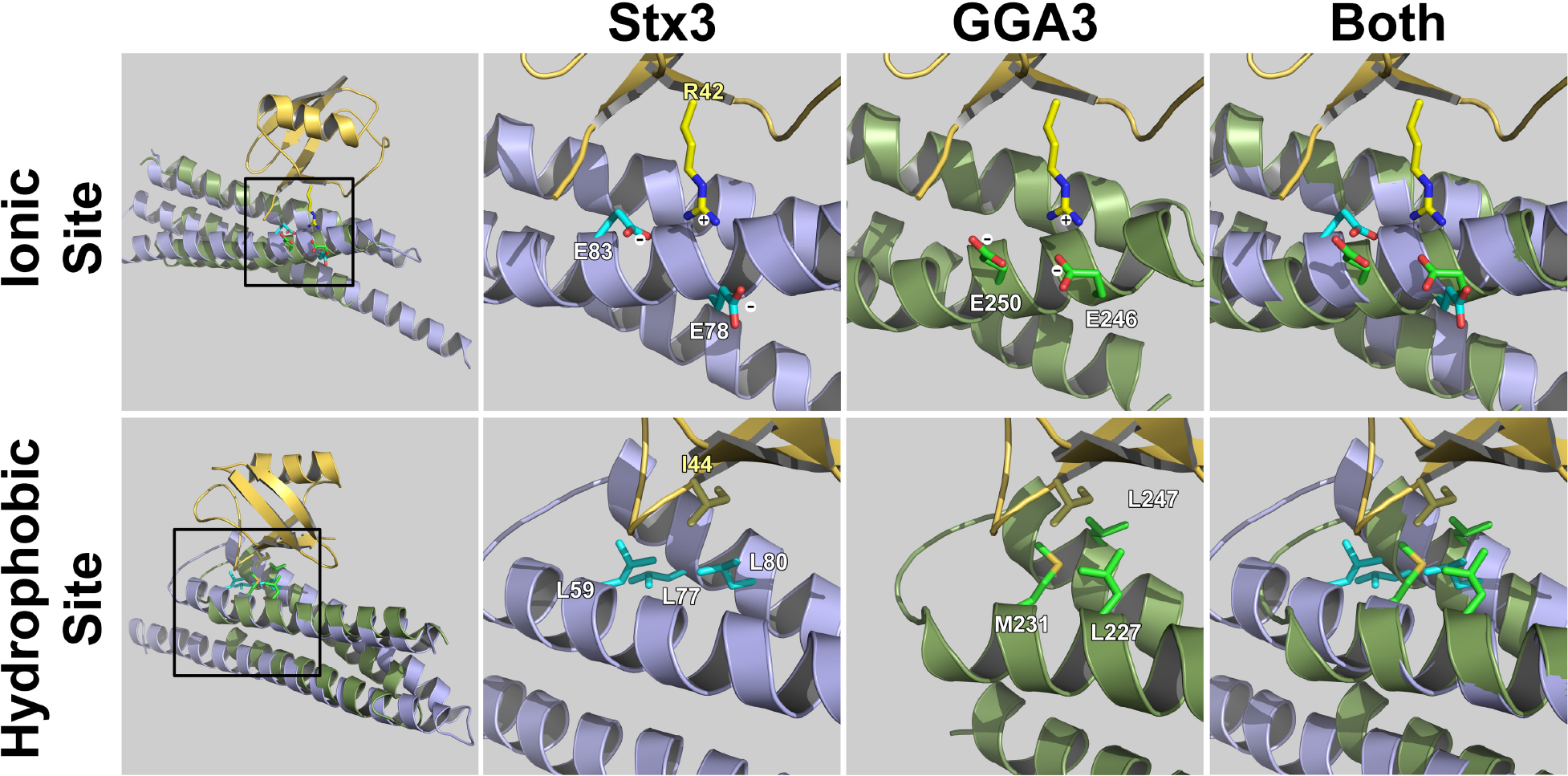
Structural modeling. **(A)** Structural model based on the X-ray structure of ubiquitin in association with site 1 of the GGA3 GAT domain [23, 24]. The Stx3 sequence was modeled into the known structure of the H_abc_ domain of the closely related Stx1A [10, 11] and fitted onto the GGA3 GAT domain. R42 of ubiquitin (yellow) is known to engage in an ionic interaction with E246 and E250 of GGA3-GAT (green). I44 of ubiquitin (yellow) is known to engage in interactions with a hydrophobic pocket formed by L227, M231 and L247 of GGA3-GAT (green). Stx3 residues (blue) that correspond most closely to these residues are E78 and E83 for the ionic site and L59, L77 and L80 for the hydrophobic site.

Due to the lack of similarity between Stx3 and GGA3 at the primary sequence level (Fig. 3A) these predictions would have been difficult or impossible to make. However, we note that E78 and E83 of Stx3 and E246 and E250 of GGA3 can be aligned closely with each other (Fig. 3A). Sequence alignment of Stx3 orthologs from numerous species indicates that all of the residues that may potentially interact with ubiquitin are highly conserved (Fig. 3B).

### Mutational analysis

Based on this model, we decided to mutate residues L77, E78, L80, and E83 of Stx3 to alanine residues and test any effects on the ability to interact with ubiquitin chains. GST fusion proteins with the cytoplasmic domain of Stx3 containing either double leucine (L77A/L80A) or double glutamate (E79A/E83A) mutations were generated. Introducing all four mutations simultaneously resulted in an insoluble GST fusion protein that could not be analyzed.

The ability of the mutants to bind to K63-linked polyubiquitin chains was assessed using the same assay as in Figure 2. Introducing the E79A/E83A mutations had no discernible effect on polyubiquitin binding as compared to wild-type Stx3 (Fig. 5). Introducing the L77A/L80A mutations had only a seemingly minor effect in that it eliminated a very weak interaction with K63-linked Ubi_2_ (Fig. 5). Given that similar mutagenesis experiments with GGA GAT domains frequently lead only to minor disruption of ubiquitin binding [19, 21, 23], however, these results may not be surprising. First, the interactions between the GAT domain and ubiquitin involve numerous contacts with multiple residues. Second, the fact that the GAT domain has two separate ubiquitin binding sites suggests that mutations of one site alone will have little effect on overall ubiquitin binding, especially for the binding of polyubiquitin chains. We predict that polyubiquitin chains wrap around the entire surface of the GAT domain, and by analogy also the H_abc_ domain of syntaxins, and engage in numerous contacts that are difficult to completely disrupt by mutagenesis. Such a binding mode may also explain why similar structures (GAT and H_abc_ domains) could bind to polyubiquitin even in the absence of highly conserved primary sequence similarity. In this regard, it is interesting that the L77A/L80A mutations in Stx3 appear to disrupt only the binding to ubiquitin dimers, which may suggest that longer ubiquitin chains can compensate by interacting with additional residues simultaneously.

**Figure 5.**
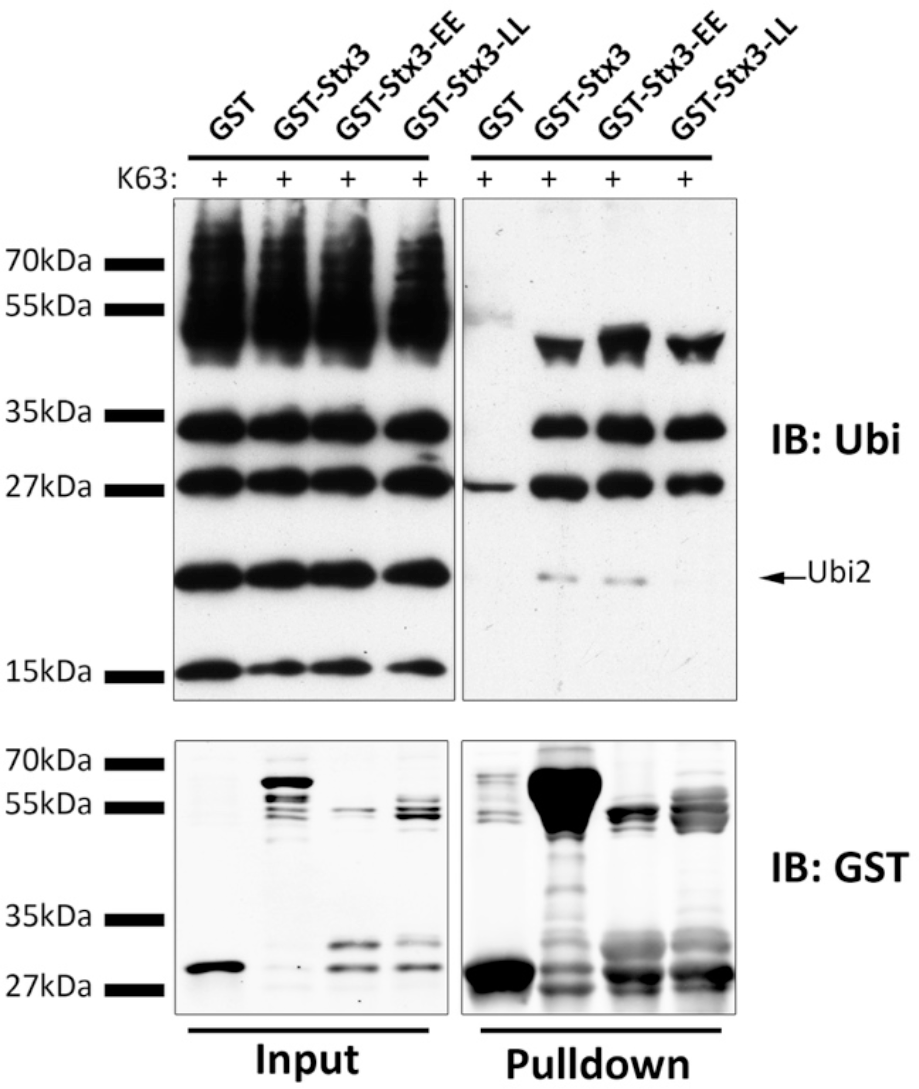
Binding mutant Stx3 to K63-linked polyubiquitin chains. Purified GST fusion protein of the wild-type Stx3 cytoplasmic region (1-265) or of mutants containing either L77A/L80A (LL) or E79A/E83A (EE) mutations were incubated with K63-linked polyubiquitin chains (1-7 ubiquitin molecules in length) followed by glutathione sepharose pull-down and IB using anti-ubiquitin and anti-GST antibodies.

## Discussion

This study illuminates a novel characteristic of syntaxins, ubiquitin binding. This is a surprising finding because, to our knowledge, ubiquitin-binding has not previously been reported for any SNARE protein, nor has there been any indication that ubiquitin-binding could affect SNARE-mediated membrane fusion events. On the other hand, the fact that the 3D structures of the GAT and H_abc_ domains are highly similar, and the fact that critical hydrophobic and ionic residues known to mediate ubiquitin-binding of the GAT domain have equivalent residues in the H_abc_ domain of Stx3 (Fig. 4) makes it plausible that both domains share a similar function.

Besides in GGA proteins, GAT domains are also present in the more distantly related proteins TOM1 and TOM1-L1, both of which also bind to ubiquitin [35]. The degree of primary sequence similarity among these GAT domains is low but the overall structures of these 3-helix bundles are highly conserved. The finding that H_abc_ domains of syntaxins not only share the same fold with GAT domains but also function in ubiquitin-binding strongly suggests that all of these domains share a common ancestry.

While the purpose of the ability of syntaxins to bind to K63-linked polyubiquitin chains remains to be elucidated, several possibilities can be envisioned. Binding of K63-linked polyubiquitin chains to the H_abc_ domain of a syntaxin may interfere with the ability of that syntaxin to bind to other regulatory proteins that are known to interact with the H_abc_ domain such as members of the munc13, synaptotagmin and munc18 families of SNARE regulators. Thereby K63-linked polyubiquitin chains, possibly attached to specific regulatory proteins, may regulate SNARE function and therefore membrane fusion in certain vesicle trafficking pathways. Another possibility is that binding of K63-linked polyubiquitin chains to the H_abc_ domain of a syntaxin would interfere with the ability of the H_abc_ domain to engage in an intramolecular interaction with the SNARE domain of that syntaxin. This would result in a “constitutively open” conformation of that syntaxin and, again, may regulate membrane fusion functions. Another possibility emerges from our recent finding that Stx3 can undergo ubiquitination at a cluster of lysine residues located between its SNARE domain and transmembrane domain [34]. It may be possible that the H_abc_ domain could engage in an intramolecular interaction with this covalently attached ubiquitin thereby locking such modified Stx3 in a “constitutively closed” conformation. Finally, it is possible that binding of K63-linked polyubiquitin chains to the H_abc_ domain of Stx3 may be unrelated to a function in membrane fusion but rather relates to a different function. Such a possibility may be supported by our recent finding that Stx3 that is covalently ubiquitinated at the lysine cluster proximal to its transmembrane domain enters the endosomal pathway, intraluminal vesicles of MVBs, and is eventually excreted with exosomes [34]. We reported that a non-ubiquitinatable mutant of Stx3 (termed Stx3-5R) is not only unable to enter the MVB/exosomal pathway but also interferes with the recruitment of a specific apical exosomal cargo protein, the orphan G-protein coupled receptor GPRC5B, into this pathway. This suggested that Stx3 normally plays a role in cargo recruitment in a fashion that is dependent on its ability to be ubiquitinated. Interestingly, the Stx3-5R mutant was also found to disrupt the secretion of human cytomegalovirus (hCMV) virions, a result that - combined with other findings - suggests that hCMV exploits the MVB/exosomal pathways for virion production and secretion [34]. In this regard, Stx3 bears striking similarities to GGA proteins. Both contain a similarly structured ubiquitin-binding domain, both undergo ubiquitination themselves, and both are involved in recruitment of membrane proteins into the MVB pathway. In the case of GGA proteins, this sorting function requires cargo proteins to be tagged with K63-linked polyubiquitin chains [31, 32].

Altogether, these results suggest that syntaxins contain a ubiquitin binding domain similar to the GAT domain. The implications of this finding are yet to be elucidated but may relate to the regulation of membrane fusion functions and/or point towards a novel function of syntaxins in the sorting of membrane proteins.

## Materials and Methods

### Plasmid construction

The cytoplasmic region of rat Stx3 (1-265) and the N-terminal region of Stx3 (1-146), respectively, were cloned into pGEX-4T-3 (GE Healthcare Life Sciences). pGEX-4T-2-GGA3-GAT plasmid was a kind gift of Kazuhisa Nakayama (Graduate School of Pharmaceutical Sciences, Kyoto University). Site-directed mutagenesis (QuickChange II, Agilent Technologies) was employed to generate the open-conformation mutant (LE165/166AA) in the Stx3 (1-265) plasmid.

### Protein expression and purification

GST-fusion protein plasmids were transformed into *E. coli* Rosetta 2 (EMD Millipore) competent cells. When the cultures reached an OD of 0.6, IPTG was added to induce expression of GST-protein. Cells were pelleted and lysed in a buffer containing 10 mM Tris-HCl, pH 8.0, 150 mM NaCl with lysozyme, RNase, DNase, Triton X-100. DTT, 5 mM EDTA, PMSF, and a protease inhibitor cocktail were subsequently added. CL2B Sepharose beads (GE Healthcare Life Sciences) were used to pre-clear the lysates, followed by incubation with glutathione-coated agarose overnight at 4°C with rotating. Beads were washed four times and eluted with glutathione. Eluate content was analyzed by SDS-PAGE. Eluates were then dialyzed into 50 mM Tris-HCl, pH 8.0, 150 mM NaCl, 1 mM EDTA and the protein concentration determined using a Nanodrop spectrophotometer (ThermoScientific).

### Mono-ubiquitin binding assay

Purified GST-fusion proteins were incubated in Buffer A (25 mM HEPES-KOH, pH 7.4, 125 mM potassium acetate, 2.5 mM magnesium acetate, 5 mM EGTA) containing 1% fetal bovine serum with ubiquitin-coated agarose beads (Sigma-Aldrich) for 2 hours at room temperature. Beads were washed three times with Buffer A containing 0.005% Tween-20, re-suspended in SDS-PAGE sample buffer, boiled, separated on SDS-PAGE, and transferred to nitrocellulose membrane. Membranes were probed with a polyclonal goat anti-GST antibody (GE Healthcare Life Sciences) and a donkey anti-goat IgG HRP-conjugated secondary antibody (Jackson Immunoresearch).

### Poly-ubiquitin binding assay

GST-fusion proteins were incubated in Buffer A containing FBS (25 mM HEPES-KOH, pH 7.4, 125 mM potassium acetate, 2.5 mM magnesium acetate, 5 mM EGTA, 1% FBS) while rotating at room temperature with 8 μg of a K48-linked or K63-linked mixture of polyubiquitin chains of 2-7 ubiquitins in length (Boston Biochem). After one hour, glutathione-coated agarose beads were added to the sample and incubated for one hour. Beads were washed three times with Buffer A containing 0.005% Tween-20, re-suspended in SDS-PAGE sample buffer and treated as above. Membranes were boiled for 10 minutes in H_2_O prior to blocking in 5% dry milk in TBST before being probed with mouse anti-ubiquitin antibody P4D1 (Santa Cruz Biotechnology).

### Surface plasmon resonance measurements

All SPR measurements were performed on a Biacore 2000 instrument. All binding assays were performed at room temperature in HBS-EP buffer (10 mM HEPES, pH 7.4, 150 mM NaCl, 3 mM EDTA, 0.005% Tween-20). GST-fusion proteins were captured to a CM5 Sensor Chip (GE Healthcare Life Sciences) by using a GST capture kit (GE Healthcare Life Sciences) according to the manufacturer’s instructions. Purified bovine ubiquitin was from Sigma-Aldrich (U6253).

## Acknowledgments

This work was supported by grants from the NIH (DK095248, GM66785), and the California Cancer Research Coordinating Committee to T.W., and a Postdoctoral Fellowship from the Spanish Ministry of Education and Science to E.R.

## References

1. Rothman JE. The principle of membrane fusion in the cell (nobel lecture). Angewandte Chemie. 2014;53(47):12676-94. doi: 10.1002/anie.201402380. PubMed PMID: 25087728.

2. Wickner W, Schekman R. Membrane fusion. Nature structural & molecular biology. 2008;15(7):658-64. PubMed PMID: 18618939; PubMed Central PMCID: PMC2488960.

3. Weimbs T, Low SH, Chapin SJ, Mostov KE, Bucher P, Hofmann K. A conserved domain is present in different families of vesicular fusion proteins: a new superfamily. Proceedings of the National Academy of Sciences of the United States of America. 1997;94(7):3046-51. Epub 1997/04/01. PubMed PMID: 9096343; PubMed Central PMCID: PMC20319.

4. Weimbs T, Mostov K, Low SH, Hofmann K. A model for structural similarity between different SNARE complexes based on sequence relationships. Trends in cell biology. 1998;8(7):260-2. PubMed PMID: 9714596.

5. Sudhof TC. The molecular machinery of neurotransmitter release (nobel lecture). Angewandte Chemie. 2014;53(47):12696-717. doi: 10.1002/anie.201406359. PubMed PMID: 25339369.

6. Fasshauer D, Sutton RB, Brunger AT, Jahn R. Conserved structural features of the synaptic fusion complex: SNARE proteins reclassified as Q- and R-SNAREs. Proceedings of the National Academy of Sciences of the United States of America. 1998;95(26):15781-6. PubMed PMID: 9861047.

7. Brunger AT. Structure and function of SNARE and SNARE-interacting proteins. Quarterly reviews of biophysics. 2005;38(1):1-47. Epub 2005/12/13. doi: 10.1017/S0033583505004051. PubMed PMID: 16336742.

8. Jahn R, Fasshauer D. Molecular machines governing exocytosis of synaptic vesicles. Nature. 2012;490(7419):201-7. doi: 10.1038/nature11320. PubMed PMID: 23060190; PubMed Central PMCID: PMCPMC4461657.

9. Han J, Pluhackova K, Bockmann RA. The Multifaceted Role of SNARE Proteins in Membrane Fusion. Front Physiol. 2017;8:5. doi: 10.3389/fphys.2017.00005. PubMed PMID: 28163686; PubMed Central PMCID: PMCPMC5247469.

10. Lerman JC, Robblee J, Fairman R, Hughson FM. Structural analysis of the neuronal SNARE protein syntaxin-1A. Biochemistry. 2000;39(29):8470-9. PubMed PMID: 10913252.

11. Fernandez I, Ubach J, Dulubova I, Zhang X, Sudhof TC, Rizo J. Three-dimensional structure of an evolutionarily conserved N-terminal domain of syntaxin 1A. Cell. 1998;94(6):841-9. PubMed PMID: 9753330.

12. Misura KM, Bock JB, Gonzalez LC, Jr., Scheller RH, Weis WI. Three-dimensional structure of the amino-terminal domain of syntaxin 6, a SNAP-25 C homolog. Proceedings of the National Academy of Sciences of the United States of America. 2002;99(14):9184-9. doi: 10.1073/pnas.132274599. PubMed PMID: 12082176; PubMed Central PMCID: PMCPMC123115.

13. Munson M, Chen X, Cocina AE, Schultz SM, Hughson FM. Interactions within the yeast t-SNARE Sso1p that control SNARE complex assembly. Nat Struct Biol. 2000;7(10):894-902. doi: 10.1038/79659. PubMed PMID: 11017200.

14. Crystal structure of syntaxin 10 from Homo sapiens (PDB ID: 4DND) [Internet]. 2012. Available from: http://www.rcsb.org/pdb/explore.do?structureId=4dnd.

15. Dulubova I, Yamaguchi T, Wang Y, Sudhof TC, Rizo J. Vam3p structure reveals conserved and divergent properties of syntaxins. Nat Struct Biol. 2001;8(3):258-64. doi: 10.1038/85012. PubMed PMID: 11224573.

16. Dietrich LE, Boeddinghaus C, LaGrassa TJ, Ungermann C. Control of eukaryotic membrane fusion by N-terminal domains of SNARE proteins. Biochimica et biophysica acta. 2003;1641(2-3):111-9. PubMed PMID: 12914952.

17. Dulubova I, Sugita S, Hill S, Hosaka M, Fernandez I, Sudhof TC, et al. A conformational switch in syntaxin during exocytosis: role of munc18. Embo J. 1999;18(16):4372-82. doi: 10.1093/emboj/18.16.4372. PubMed PMID: 10449403; PubMed Central PMCID: PMCPMC1171512.

18. Suer S, Misra S, Saidi LF, Hurley JH. Structure of the GAT domain of human GGA1: a syntaxin amino-terminal domain fold in an endosomal trafficking adaptor. Proceedings of the National Academy of Sciences of the United States of America. 2003;100(8):4451-6. Epub 2003/04/02. doi: 10.1073/pnas.0831133100. PubMed PMID: 12668765; PubMed Central PMCID: PMC404691.

19. Shiba Y, Katoh Y, Shiba T, Yoshino K, Takatsu H, Kobayashi H, et al. GAT (GGA and Tom1) domain responsible for ubiquitin binding and ubiquitination. The Journal of biological chemistry. 2004;279(8):7105-11. doi: 10.1074/jbc.M311702200. PubMed PMID: 14660606.

20. Puertollano R, Bonifacino JS. Interactions of GGA3 with the ubiquitin sorting machinery. Nature cell biology. 2004;6(3):244-51. doi: 10.1038/ncb1106. PubMed PMID: 15039775.

21. Scott PM, Bilodeau PS, Zhdankina O, Winistorfer SC, Hauglund MJ, Allaman MM, et al. GGA proteins bind ubiquitin to facilitate sorting at the trans-Golgi network. Nature cell biology. 2004;6(3):252-9. doi: 10.1038/ncb1107. PubMed PMID: 15039776.

22. Bilodeau PS, Winistorfer SC, Allaman MM, Surendhran K, Kearney WR, Robertson AD, et al. The GAT domains of clathrin-associated GGA proteins have two ubiquitin binding motifs. J Biol Chem. 2004;279(52):54808-16. doi: 10.1074/jbc.M406654200. PubMed PMID: 15494413; PubMed Central PMCID: PMCPMC2911622.

23. Prag G, Lee S, Mattera R, Arighi CN, Beach BM, Bonifacino JS, et al. Structural mechanism for ubiquitinated-cargo recognition by the Golgi-localized, gamma-ear-containing, ADP-ribosylation-factor-binding proteins. Proceedings of the National Academy of Sciences of the United States of America. 2005;102(7):2334-9. doi: 10.1073/pnas.0500118102. PubMed PMID: 15701688; PubMed Central PMCID: PMCPMC549010.

24. Kawasaki M, Shiba T, Shiba Y, Yamaguchi Y, Matsugaki N, Igarashi N, et al. Molecular mechanism of ubiquitin recognition by GGA3 GAT domain. Genes to cells: devoted to molecular & cellular mechanisms. 2005;10(7):639-54. doi: 10.1111/j.1365-2443.2005.00865.x. PubMed PMID: 15966896.

25. Pelham HR. Membrane traffic: GGAs sort ubiquitin. Current biology: CB. 2004;14(9):R357-9. doi: 10.1016/j.cub.2004.04.027. PubMed PMID: 15120092.

26. Mukhopadhyay D, Riezman H. Proteasome-independent functions of ubiquitin in endocytosis and signaling. Science. 2007;315(5809):201-5. Epub 2007/01/16. doi: 10.1126/science.1127085. PubMed PMID: 17218518.

27. Hicke L. A new ticket for entry into budding vesicles-ubiquitin. Cell. 2001;106(5):527-30. Epub 2001/09/12. PubMed PMID: 11551499.

28. Hessvik NP, Llorente A. Current knowledge on exosome biogenesis and release. Cell Mol Life Sci. 2017. doi: 10.1007/s00018-017-2595-9. PubMed PMID: 28733901.

29. Hurley JH, Lee S, Prag G. Ubiquitin-binding domains. Biochem J. 2006;399(3):361-72. doi: 10.1042/BJ20061138. PubMed PMID: 17034365; PubMed Central PMCID: PMCPMC1615911.

30. Hochstrasser M. Origin and function of ubiquitin-like proteins. Nature. 2009;458(7237):422-9. doi: 10.1038/nature07958. PubMed PMID: 19325621; PubMed Central PMCID: PMCPMC2819001.

31. Lauwers E, Jacob C, Andre B. K63-linked ubiquitin chains as a specific signal for protein sorting into the multivesicular body pathway. The Journal of cell biology. 2009;185(3):493-502. doi: 10.1083/jcb.200810114. PubMed PMID: 19398763; PubMed Central PMCID: PMCPMC2700384.

32. Deng Y, Guo Y, Watson H, Au WC, Shakoury-Elizeh M, Basrai MA, et al. Gga2 mediates sequential ubiquitin-independent and ubiquitin-dependent steps in the trafficking of ARN1 from the trans-Golgi network to the vacuole. The Journal of biological chemistry. 2009;284(35):23830-41. doi: 10.1074/jbc.M109.030015. PubMed PMID: 19574226; PubMed Central PMCID: PMCPMC2749155.

33. Santonico E, Mattioni A, Panni S, Belleudi F, Mattei M, Torrisi MR, et al. RNF11 is a GGA protein cargo and acts as a molecular adaptor for GGA3 ubiquitination mediated by Itch. Oncogene. 2015;34(26):3377-90. doi: 10.1038/onc.2014.256. PubMed PMID: 25195858.

34. Giovannone AJ, Reales E, Bhattaram P, Fraile-Ramos A, Weimbs T. Monoubiquitination of syntaxin 3 leads to retrieval from the basolateral plasma membrane and facilitates cargo recruitment to exosomes. Mol Biol Cell. 2017;28(21):2843-53. doi: 10.1091/mbc.E17-07-0461. PubMed PMID: 28814500; PubMed Central PMCID: PMCPMC5638587.

35. Yamakami M, Yoshimori T, Yokosawa H. Tom1, a VHS domain-containing protein, interacts with tollip, ubiquitin, and clathrin. J Biol Chem. 2003;278(52):52865-72. doi: 10.1074/jbc.M306740200. PubMed PMID: 14563850.

